# The shirker’s dilemma and the prospect of cooperation in large groups

**DOI:** 10.1101/2023.04.06.535867

**Authors:** Jorge Peña, Aviad Heifetz, Georg Nöldeke

**Affiliations:** Department of Social and Behavioral Sciences, Toulouse School of Economics; Institute for Advanced Study in Toulouse; Department of Human Behavior, Ecology and Culture, Max Planck Institute for Evolutionary Anthropology; Department of Management and Economics, Open University of Israel; Faculty of Business and Economics, University of Basel

**Keywords:** replicator dynamics, evolutionary game theory, collective action, cooperation, group size

## Abstract

Cooperation usually becomes harder to sustain as groups become larger because incentives to shirk increase with the number of potential contributors to collective action. But is this always the case? Here we study a binary-action cooperative dilemma where a public good is provided as long as not more than a given number of players shirk from a costly cooperative task. We find that at the stable polymorphic equilibrium, which exists when the cost of cooperation is low enough, the probability of cooperating increases with group size and reaches a limit of one when the group size tends to infinity. Nevertheless, increasing the group size may increase or decrease the probability that the public good is provided at such an equilibrium, depending on the cost value. We also prove that the expected payoff to individuals at the stable equilibrium (i.e., their fitness) decreases with group size. For low enough costs of cooperation, both the probability of provision of the public good and the expected payoff converge to positive values in the limit of large group sizes. However, we also find that the basin of attraction of the stable polymorphic equilibrium is a decreasing function of group size and shrinks to zero in the limit of very large groups. Overall, we demonstrate non-trivial comparative statics with respect to group size in an otherwise simple collective action problem.

## 1 Introduction

Living in groups, together with the possibilities and opportunities for conflict and cooperation entailed by communal life, is a widespread phenomenon in the natural world (Krause and Ruxton, 2002). Among the many factors affecting the evolution of cooperation in social groups, including relatedness and repeated interactions, group size has received considerable attention. In particular, how the strength of selection on cooperative behavior might increase or decrease with group size has been a recurrent question in behavioral ecology and evolutionary biology (Beauchamp and Ruxton, 2003; MacNulty et al., 2012; Shen et al., 2014; Powers and Lehmann, 2017; Peña and Nöldeke, 2018). This question is paralleled in the social sciences by the question of how individual incentives to cooperate in a collective action problem are affected by group size (Olson, 1965; Chamberlin, 1974; Palfrey and Rosenthal, 1984; Oliver and Marwell, 1988). Although there is broad consensus that the effect of group size on cooperation is typically negative, with cooperation becoming more difficult to evolve or sustain as groups become larger (Olson, 1965; Boyd and Richerson, 1988), positive group size effects have been also documented in empirical studies (Isaac et al., 1994; Powers and Lehmann, 2017) and demonstrated in theoretical research (Esteban and Ray, 2001; Cheikbossian and Fayat, 2018; Peña and Nöldeke, 2018).

To fix ideas, consider a simple game-theoretic model of social interactions: the “volunteer’s dilemma” (Diekmann, 1985). In this game, *n* players simultaneously become aware of a costly task requiring at least one volunteer. If at least one of the players volunteers to pay the cost *c* of performing the task, a good of normalized value of one is created and enjoyed by all players. If nobody volunteers, the task is left undone, and no public good is created. A more general version of this game (which, for simplicity, we will also refer to as a “volunteer’s dilemma”) considers the case where the cooperation of at least *θ* volunteers is needed for the collective good to be created, for 1 ≤ *θ* ≤ *n* (Palfrey and Rosenthal, 1984). Such games have become influential in evolutionary biology, where they have been used to model a wide array of collective action problems ranging from the secretion of extracellular compounds in microbes and the construction of collective stalks in social amoeba (Archetti, 2009) to leadership in animal societies (Shen et al., 2010; Smith et al., 2016), confrontational scavenging in hominins (Bickerton and Szathmáry, 2011), and the costly punishment of free riders in humans (Raihani and Bshary, 2011; Przepiorka and Diekmann, 2013; Schoenmakers et al., 2014).

The volunteer’s dilemma has provided theoretical underpinnings to the idea that larger groups are less conducive to cooperation. Indeed, it is well known that when only one volunteer is required (*θ* = 1) both the proportion of volunteers and the probability that the collective action is successful decreases with group size (Archetti, 2009). More recently, Nöldeke and Peña (2020) have shown that these two results extend to 1 *< θ < n* for the best symmetric Nash equilibrium of the game (i.e., the equilibrium sustaining the highest probability of cooperation among the symmetric Nash equilibria) while additionally proving that the expected payoff at such an equilibrium is also a decreasing function of group size. Together, these results establish for the volunteer’s dilemma that there are negative group-size effects on three different quantities: the proportion of volunteers, the probability that the collective action is successful, and the expected payoff at equilibrium. This casts doubts on the extent to which large-scale cooperation modelled after the volunteer’s dilemma can be sustained without the presence of additional cooperation-enhancing mechanisms and raises the question of whether such negative group-size effects are a general theoretical feature of a wide class of collective action problems or a peculiarity of the volunteer’s dilemma.

Here, we consider the group-size effects of a related cooperative dilemma, which we will call the “shirker’s dilemma”. As in the volunteer’s dilemma, *n* players simultaneously become aware of a costly collective task. Yet, while in the volunteer’s dilemma the collective task is successful (and a public good is produced) *if at least* a fixed number *θ* of players volunteer, in the shirker’s dilemma the collective task is successful *if at most* a fixed number *ζ* of players shirk from volunteering. Obviously, for a given group size *n* and cost of volunteering *c*, the shirker’s dilemma and the volunteer’s dilemma are equivalent: a shirker’s dilemma with threshold *ζ* = *n* − *θ* is nothing but a volunteer’s dilemma with threshold *θ*. However, and importantly, the two games capture different consequences of an increase in group size: While in the volunteer’s dilemma the number of required volunteers is a constant independent of group size, in the shirker’s dilemma the number of required volunteers *θ* increases with group size so as to keep the threshold maximum number of shirkers *ζ* fixed. Thus, for large group sizes (and relatively small thresholds), collective action is successful in the volunteer’s dilemma if *some* individuals cooperate. By contrast, in the shirker’s dilemma collective action is successful if *few* individuals shirk, and hence that *most* individuals cooperate. As will be shown, the different payoff structure of the shirker’s dilemma translates into different comparative statics with respect to group size and into a different answer to the question of whether or not larger groups are less conducive to cooperation.

The shirker’s dilemma is relevant for situations where the (potentially costly) cooperation of all or most individuals is required to produce a collective good. Such scenarios may arise in cases traditionally conceptualized as volunteer’s dilemmas. For instance, Archetti (2009) lists as an example of a volunteer’s dilemma the case of the amoeba *Dictyostelium discoideum* that, when facing starvation, differentiates into a ball of reproductive spores (shirkers) and a sterile stalk (volunteers). However, it can be argued that what is needed for collective action to be successful in this case is not that enough cells become part of the stalk, but that there is a cap on the number of reproductive cells that must be maintained at the top. Likewise, Raihani and Bshary (2011) argue that the punishment of free riders in an *n*-player social dilemma is best described by assuming that the benefit of punishment is a step function of the amount of punishment (e.g., the number of punishers in a group). However, for many social situations, it might be more plausible that the inflection point of such a step function (and hence the success of punishment as a collective action) is determined more by the maximum number of allowed second-order free riders (i.e., individuals that will refrain from punishing free riders) than by the minimum number of punishers. A maximal allowable number of shirkers instead of a minimal required number of volunteers— and hence the payoff structure of the shirker’s dilemma in contrast to that of a volunteer’s dilemma—may also be relevant in other human interactions. For example, a hacker may scan the computer systems of target organizations for vulnerable entry points left open by employees who compromised computer security instructions. By itself, a particular entry point does not yet guarantee successful hacking of the organization’s website or data for ransom purposes. Therefore, the hacker may decide to attack all vulnerable access points simultaneously only if their number exceeds a certain threshold and otherwise continue its search for a more vulnerable organization. In all of these situations, the shirker’s dilemma might better represent the social dilemma at hand than the volunteer’s dilemma.

Our analysis of the group-size effects on the shirker’s dilemma builds upon previous insights derived in Peña and Nöldeke (2018) for the general case of binary-action symmetric *n*-player games and in Nöldeke and Peña (2020) for the particular case of the volunteer’s dilemma. The question asked in Nöldeke and Peña (2020) was: what are the consequences of larger group sizes in a collective action that is successful if and only if there are enough contributors? Here, we ask the dual question, namely: What are the consequences of larger group sizes in a collective action that is successful if and only if there are not too many shirkers? In particular, we are interested in the three group-size effects analyzed by Nöldeke and Peña (2020) for the volunteer’s dilemma, namely: (i) the effect of group size on the probability that individuals cooperate at equilibrium, (ii) the effect of group size on the expected payoff of individuals, and (iii) the effect of group size on the probability that collective action is successful at equilibrium. Finally, we also investigate what happens to these quantities as the groups are made arbitrarily large, that is, in the limit of infinitely large groups.

## 2 Model

### 2.1 The shirker’s dilemma

Our model is a particular case of a multi-player matrix game (Broom et al., 1997) with two pure strategies (for related models see, e.g., Bach et al. 2006; Archetti 2009; Peña et al. 2014; Broom et al. 2019; Peña and Nöldeke 2023). Specifically, *n >* 2 players face a task to be performed that requires collective action for its success. Each player *i* ∈ {1, …, *n*} can either “volunteer” (or “cooperate”) at a cost *c* ∈ (0, 1) or, alternatively, “shirk” from the task (or “defect”). All players, irrespective of their strategy, enjoy an additional payoff benefit normalized to one (i.e., a public good of value one is provided) if at most *ζ* ≥ 1 players shirk. Throughout the following, we take *ζ* and *c* as given and study the impact of group size *n* on equilibrium behavior. In doing so, we restrict attention to group sizes *n > ζ* + 1, thereby excluding the trivial case in which the benefit arises even if all players shirk (*n* = *ζ*) and the well-understood case (Diekmann, 1985) in which exactly one volunteer is required for the benefit to arise (*n* = *ζ* + 1).

As noted in the Introduction, for given *n* our game corresponds to a volunteer’s dilemma with *θ* = *n* − *ζ* as the minimal number of volunteers. Nevertheless, we prefer the name “shirker’s dilemma” as it draws attention to the essential feature of our analysis, namely that we study the case in which the maximal number of shirkers (rather than the minimal number of volunteers) compatible with the provision of the benefit is considered fixed.

### 2.2 Replicator dynamic and pivot probability

We assume that the population is infinitely large and comprised of at most two types of individuals: “shirkers” (or “defectors”) and “volunteers” (or “cooperators”). Denoting by *q* the proportion of shirkers in the population, and assuming that groups of size *n* are matched to play the game uniformly at random, the probability that there are exactly *k* shirkers in a group is given by

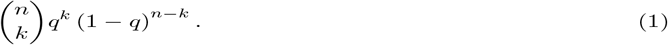

Letting

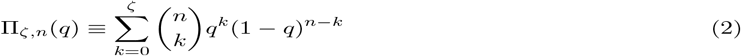

denote the probability that there are at most *ζ* shirkers in a group, the expected payoff to a shirker and to a volunteer can then be written as

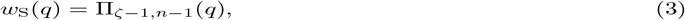

and

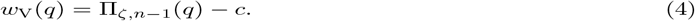

Indeed, a shirker gets a payoff of one if there are not more than *ζ* − 1 shirkers among its *n* − 1 co-players, and zero otherwise, which explains Eq.(3). In contrast, a volunteer gets a benefit of one in case there are not more than *ζ* shirkers among its co-players but always pays the cost *c* of volunteering, which explains Eq.(4). Note that Eqs. (1) and (2) correspond, respectively, to the probability mass function and the cumulative distribution function of a binomial distribution with parameters *n* and *q*.

We assume that the change in the proportion of shirkers over evolutionary time is given by the continuous-time two-strategy replicator dynamic (Taylor and Jonker, 1978; Weibull, 1995; Hofbauer and Sigmund, 1998)

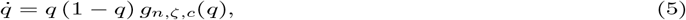

where *g*_*n,ζ,c*_(*q*) corresponds to the “gain function” (Bach et al., 2006; Peña et al., 2014), the “incentive function” (Broom and Rychtář, 2022, p. 177-178), or the “private gain function” (Peña and Nöldeke, 2023). The gain function is the difference in expected payoffs between the two pure strategies in a population where the proportion of shirkers is *q* and quantifies the selection pressure on the proportion of shirkers in the population. In our case, the gain function is given by

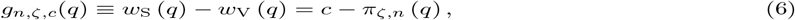

where

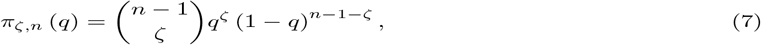

is the probability that *ζ* out of *n* − 1 groups members shirk when the proportion of shirkers in the population is equal to *q*. As a given focal player’s strategy is decisive for whether or not the collective action of the group is successful exactly when *ζ* other group members shirk, Eq.(7) is thus the probability that a change in the focal player’s strategy changes the outcome of the social interaction. This is analogous to what is known as the pivot probability in the game-theoretic analysis of voting models (Palfrey and Rosenthal, 1983; Nöldeke and Peña, 2016), i.e., the probability that the decision of a single voter will change the outcome of an election.

### 2.3 Rest points and their stability

We are interested in the asymptotically stable rest points of the replicator dynamic (5) as these indicate the effects of selection in the long run for our model. The rest points of the replicator dynamics correspond to all values of the share of shirkers *q* that nullify Eq.(5). There are two kinds of rest points. First, there are two trivial rest points *q* = 0 (“all volunteer”) and *q* = 1 (“all shirk”) at which the population is monomorphic and the type variance of the population, given by *q*(1 − *q*), is equal to zero. Second, there can be interior rest points *q* ∈ (0, 1), at which the population is polymorphic but the expected payoffs to the two strategies are equal (i.e., Π_S_ (*q*) − Π_V_ (*q*) = 0 holds), so that the gain function vanishes. Setting Eq.(6) to zero, we find that the interior rest points correspond to the solutions of the pivotality condition

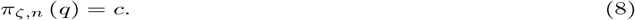

The stability of both trivial and interior rest points can be verified by checking the sign pattern of the gain function (Bukowski and Miekisz, 2004; Peña et al., 2014; Peña and Nöldeke, 2023). In particular, *q* = 0 is stable if the initial sign of the gain function is negative, *q* = 1 is stable if the final sign of the gain function is positive, and an interior rest point is stable (resp. unstable) if the gain function changes sign from positive to negative (resp. negative to positive) at the rest point. In turn, the sign pattern of the gain function (6) depends on how the pivot probability compares to the cost of volunteering as a function of the proportion of shirkers in the population. To make progress, we thus need a full characterization of the shape of the pivot probability as a function of the proportion of shirkers.

To this end, it can be verified that the pivot probability *π*_*ζ,n*_ (*q*) satisfies the following properties (see Nöldeke and Peña, 2020; the top left panel of Fig. 1 illustrates). First, the pivot probability is differentiable in *q* (differentiability). Second, *π*_*ζ,n*_ (0) = *π*_*ζ,n*_ (1) = 0 holds (end-points property). Third, the pivot probability is strictly increasing on the interval [0, *ζ/*(*n* − 1)] and strictly decreasing on the interval [*ζ/*(*n* − 1), 1] with non-zero derivative on the interiors of these intervals (unimodality). In particular, *ζ/*(*n* − 1) is the unique maximizer of *π*_*ζ,n*_ (*q*) in the interval [0, 1]. Hence,

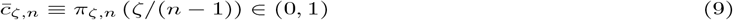

is the *critical cost* value such that (i) for 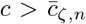 the pivotality condition (8) has no solution, (ii) for 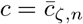 it has a unique solution, and (iii) for 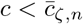 it has two solutions.

**Figure 1.**
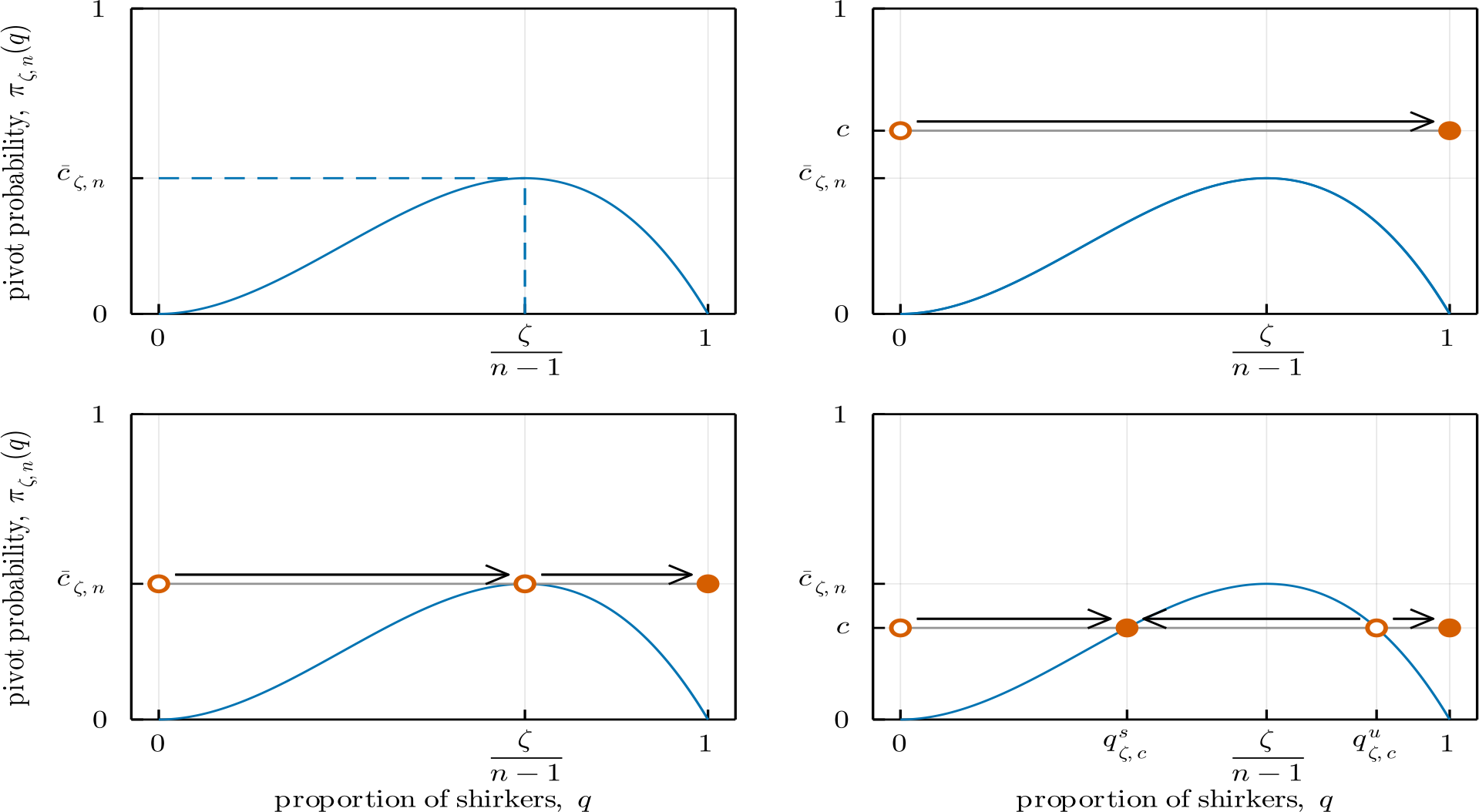
Pivot probability *π*_*ζ,n*_ (*q*) as a function of the proportion of shirkers *q*, and corresponding evolutionary dynamics, as given by Lemma 1, for *ζ* = 2 and *n* = 4. *Top left:* The pivot probability is unimodal, with maximum 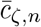 at *q* = *ζ/*(*n* − 1). Here, 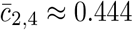, and *ζ/*(*n* − 1) = 2*/*3. *Top right:* For high costs 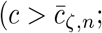 here *c* = 0.6) the replicator dynamic has no interior rest points. The trivial rest point *q* = 0 is unstable (*open circle*) while the trivial rest point *q* = 1 is stable (*full circle*). The evolutionary dynamics lead to *q* = 1 for all initial conditions (*arrow*). *Bottom left:* For a cost equal to the critical cost 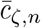, there is a unique interior rest point at *q* = *ζ/*(*n* − 1) that is unstable (*open circle*). The evolutionary dynamics lead to *q* = 1 for all initial conditions in [0, 1] − {*ζ/*(*n* − 1)} (*arrows*). *Bottom right:* For low costs (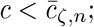 here, *c* = 0.3), the replicator dynamic features two interior rest points: the smaller 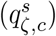 is stable (*full circle*), and the larger 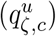 is unstable (*open circle*). The dynamics (*arrows*) lead either to 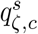 or to *q* = 1, depending on initial conditions.

The shape properties of the pivot probability allow us to fully characterize the evolutionary dynamics of the shirker’s dilemma. Indeed, it follows from the end-points property and the assumption *c >* 0 that *q* = 0 (“all volunteer”) is always an unstable rest point of the replicator dynamic (5), while *q* = 1 (“all shirk”) is always a stable rest point. Additionally, these are the only rest points when the cost of volunteering is sufficiently large (i.e., when 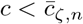 holds). When the cost is sufficiently low (i.e., when 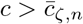 holds) the replicator dynamic has two additional rest points in the interval (0, 1), corresponding to the two solutions to the pivotality condition (8). The derivative of the gain function is negative at the smaller, while it is positive at the larger of these rest points. Hence, the gain function changes sign from positive to negative at the smaller of these rest points (which is then stable), while it changes sign from negative to positive at the larger of these rest points (which is then unstable). We collect these observations on the characterization of the replicator dynamic of the shirker’s dilemma in the following Lemma (cf. Lemma 1 in Nöldeke and Peña, 2020; the top right and bottom panels of Fig. 1 illustrate).

#### Lemma 1.

*For any ζ, n, and c, q* = 0 *is an unstable rest point and q* = 1 *is a stable rest point of the replicator dynamic. Additionally, the number, location, and stability of interior rest points of the replicator dynamic depend on how the cost of volunteering c compares to the critical cost* (9) *as follows:*

1. *If* 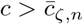, *there are no interior rest points*.
2. *If* 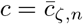, *there is a unique unstable interior rest point, namely q* = *ζ/*(*n* − 1).
3. *If* 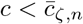, *there are two interior rest points:* 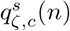, *which is stable, and* 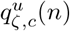, *which is unstable*. *The two interior rest points satisfy* 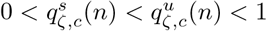, *with* 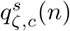 *being the unique solution to* (8) *in the interval* (0, *ζ/*(*n* − 1)), *and* 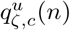 *being the unique solution to* (8) *in the interval* (*ζ/*(*n* − 1), 1).

There are two cases of interest. First, for high costs (i.e., 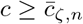) the replicator dynamic is such that the only stable rest point is *q* = 1, i.e., the population is characterized by full shirking at equilibrium. Second, for low costs (i.e., 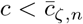, the replicator dynamic exhibit what can be called bistable coexistence (Peña et al., 2015), where an interior unstable rest point (located at 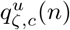 separates the basins of attraction of two stable rest points: one where there is full shirking (*q* = 1) and another one where shirkers and volunteers coexist (i.e., 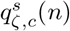).

### 2.4 Proportion of volunteers, success probability, and expected payoff

We are interested in the effects of group size on three quantities: (i) the proportion of volunteers, (ii) the probability of collective success, and (iii) the expected payoff, all evaluated at the stable rest point of the replicator dynamic sustaining the smallest amount of shirking (i.e., the largest amount of volunteering). In the following, we refer to such rest point as the *minimal rest point* and denote it by *q*_*ζ,c*_(*n*).

Depending on the particular values of *ζ, n*, and *c*, the minimal rest point can be either the trivial rest point *q* = 1 or the interior rest point 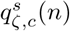. More precisely, from Lemma 1 we have

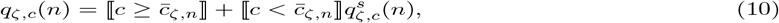

where we have used the Iverson bracket to write ⟬*P*⟭ = 1 if *P* is true and ⟬*P* ⟭= 0 if *P* is false. With this definition of *q*_*ζ,c*_(*n*) we let

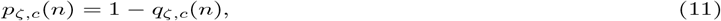

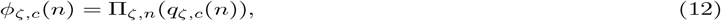

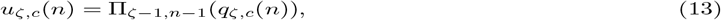

denote, respectively, the *proportion of volunteers*, the *success probability*, and the *expected payoff* at the minimal rest point. The proportion of volunteers is simply one minus the proportion of shirkers at the minimal rest point. It is thus given by Eq.(11). The success probability is the probability that the collective good is produced. This happens when there are not more than *ζ* shirkers in a group of *n* players all of whom are shirking with probability (10). It is thus given by Eq.(12). Finally, the expected payoff at the minimal rest point is most easily calculated from the perspective of a shirker (see Eq.(3)), and thus given by Eq.(13). Clearly, the equilibrium payoff can be also calculated from an ex-ante perspective, and is thus given by the success probability minus the expected cost of volunteering. This observation provides a useful equation linking the three quantities, namely

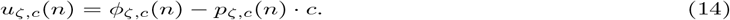

## 3 Results

### 3.1 Preliminaries: The effect of group size on the critical cost

Before proceeding to our main results, we first need to characterize how the critical cost 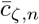 from Eq.(9) depends on group size. This is important because the critical cost determines whether the minimal rest point *q*_*ζ,c*_(*n*) from Eq.(10) features full shirking or a positive proportion of volunteers. The following result provides such a characterization and an additional result concerning the relation between the threshold *ζ* and the critical cost. The proof is in Appendix A.1. Fig. 2 illustrates.

**Figure 2.**
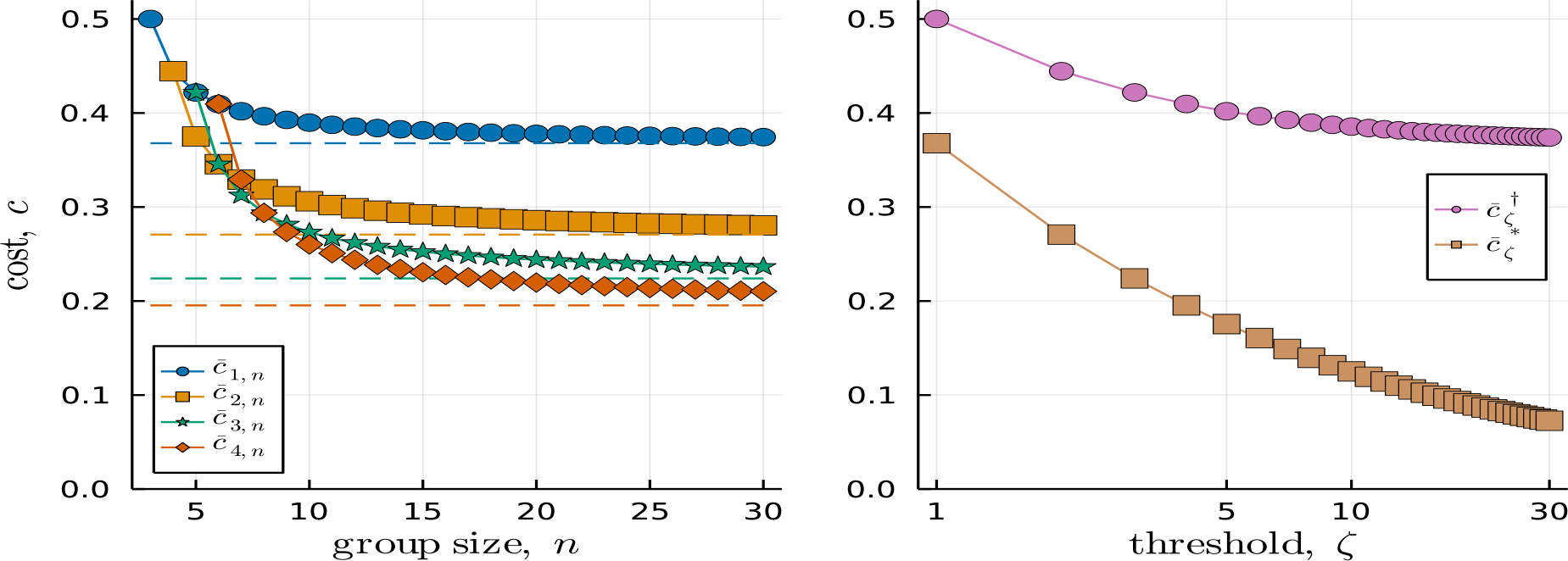
Critical cost 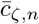, maximum critical cost 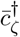, and limit critical cost 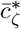. *Left:* Critical cost 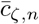(*circles*) as a function of group size *n* for *ζ* ∈ {1, 2, 3, 4}. As stated in Lemma 2, the sequence of critical costs is strictly decreasing for all *ζ* with a limit given by 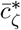(*dashed lines*). *Right:* Maximum critical cost 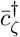(*circles*) and limit critical cost 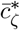(*squares*) as a function of threshold *ζ*.

#### Lemma 2.

*For any ζ, the critical cost* 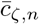 *is strictly decreasing in n with maximum*

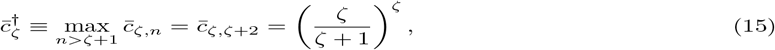

*and limit*

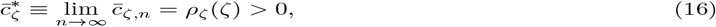

*where*

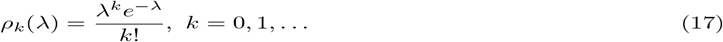

*denotes the probability mass function of a Poisson distribution with parameter λ*.

*Moreover, both* 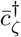*and* 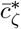 *are decreasing in ζ, with limits* 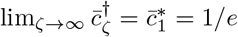, *and* 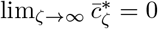.

Lemma 2 indicates a negative effect of group size on the evolution of cooperation for the shirker’s dilemma. Namely, increasing the group size decreases the critical cost value 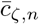, and hence increases the range of cost levels for which full shirking is the unique stable rest point. In particular, for cost values satisfying 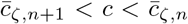, some cooperation can be sustained at equilibrium for the smaller group size *n* (i.e., *p*_*ζ,c*_(*n*) *>* 0) but not for the larger group size *n* + 1 (i.e., *p*_*ζ,c*_(*n* + 1) = 0). An analogous negative group-size effect is present in the volunteer’s dilemma (Nöldeke and Peña, 2020, Lemma 1).

An immediate consequence of Lemma 2 is that, for a given threshold *ζ* ≥ 1, the cost *c* ∈ (0, 1) can fall into one of three different regions (see right panel of Fig. 2).

1. For 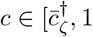, costs are so high that 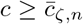 holds for all group sizes *n*. In this case, the minimal rest point is given by *q*_*ζ,c*_(*n*) = 1 for all *n*. Further, the proportion of volunteers, the success probability, and the expected payoff all reduce to

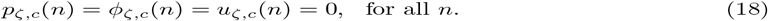
2. For 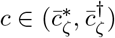, there exists a finite *critical group size* 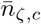, such that 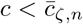holds if and only if 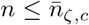 holds, while 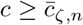 holds if and only if 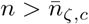 holds. In this case, the minimal rest point corresponds to the non-trivial rest point 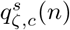 for 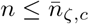, and to the trivial rest point *q* = 1 for 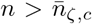. It follows that the proportion of volunteers, the success probability, and the expected payoff are all positive for group sizes smaller than or equal to the critical group size, but they all drop to zero thereafter, namely

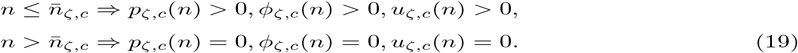
3. For 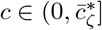, costs are sufficiently low that 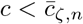 holds for all group sizes *n*. In this case, the minimal rest point corresponds to the non-trivial rest point 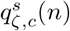, and the proportion of volunteers, the success probability, and the expected payoff are all positive for all group sizes *n*. That is, we have

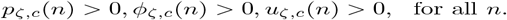

In the following, we restrict our analysis to costs satisfying 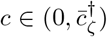 (i.e., cases 2 and 3 above), thus excluding the uninteresting case in which the proportion of volunteers, the expected payoff, and the success probability are all zero for all group sizes (i.e., case 1 above).

### 3.2 The effect of group size on the proportion of volunteers

Our first main result on group-size effects pertains to the comparative statics of the proportion of volunteers (11) at the minimal rest point (10). To derive this result, we begin by stating:

#### Lemma 3.

*For any ζ, n, and cost* 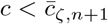, *the interior rest points of the replicator dynamic for group*

*size n and group size n* + 1 *satisfy*

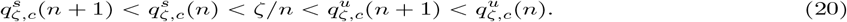

The formal proof of Lemma 3 is in Appendix A.2. Fig. 3 illustrates the underlying arguments for the inequalities 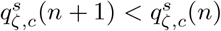 and 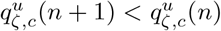. At the stable interior rest point 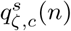, the probability that a focal group member is pivotal is *increasing* both in group size and in the proportion of shirkers. The reason that an increase in group size decreases the proportion of shirkers in the stable interior rest point is thus that the *positive* effect of an increase in group size on the pivot probability has to be compensated by a decrease in the proportion of shirkers to restore the pivotality condition. In contrast, at the unstable interior rest point 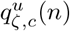, the probability that a focal group member is pivotal is *decreasing* both in group size and in the proportion of shirkers. The reason that an increase in group size decreases the proportion of shirkers in the unstable interior rest point is thus that the *negative* effect of an increase in group size on the pivot probability has to be compensated by a decrease in the proportion of shirkers to restore the pivotality condition.

**Figure 3.**
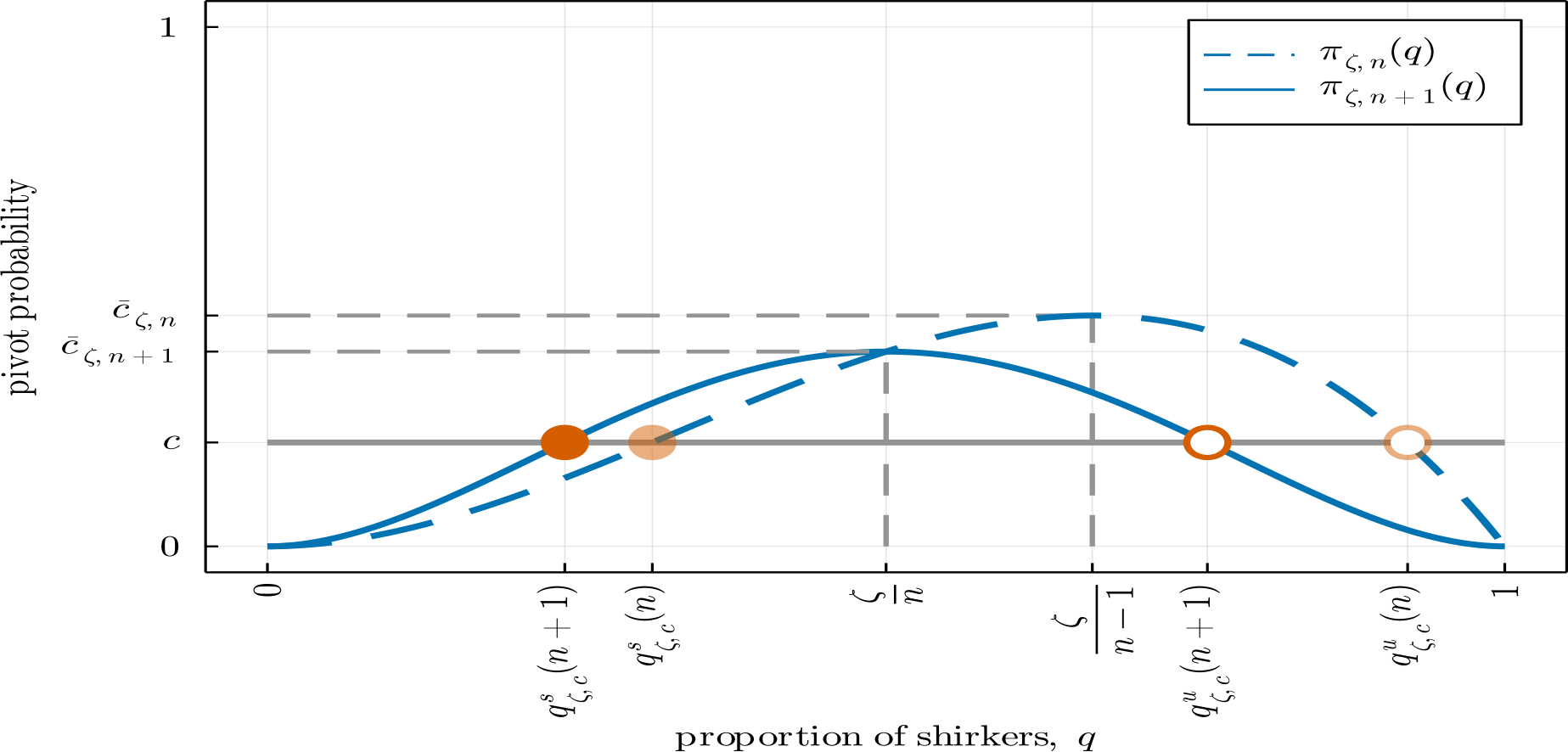
Illustration of Lemma 3 and its proof (in Appendix A.2) for *ζ* = 2, *n* = 4, and *c* = 0.2. For cost *c* lower than 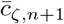, both a stable (*full circle*) and an unstable (*open circle*) rest point exist for group sizes *n* and *n* + 1, and are ordered in such a way that Eq.(20) is satisfied, with both the stable and the unstable rest points decreasing with group size.

Rewriting the first inequality in (20) in terms of the proportion of volunteers, we obtain that

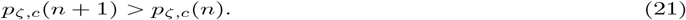

holds for any cost of volunteering 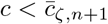. Combining this observation with Lemmas 1 and 2 directly implies the following characterization of the group-size effect on the proportion of volunteers at the minimal rest point. The left panel of Fig. 4 and Fig. 5 illustrate.

**Figure 4.**
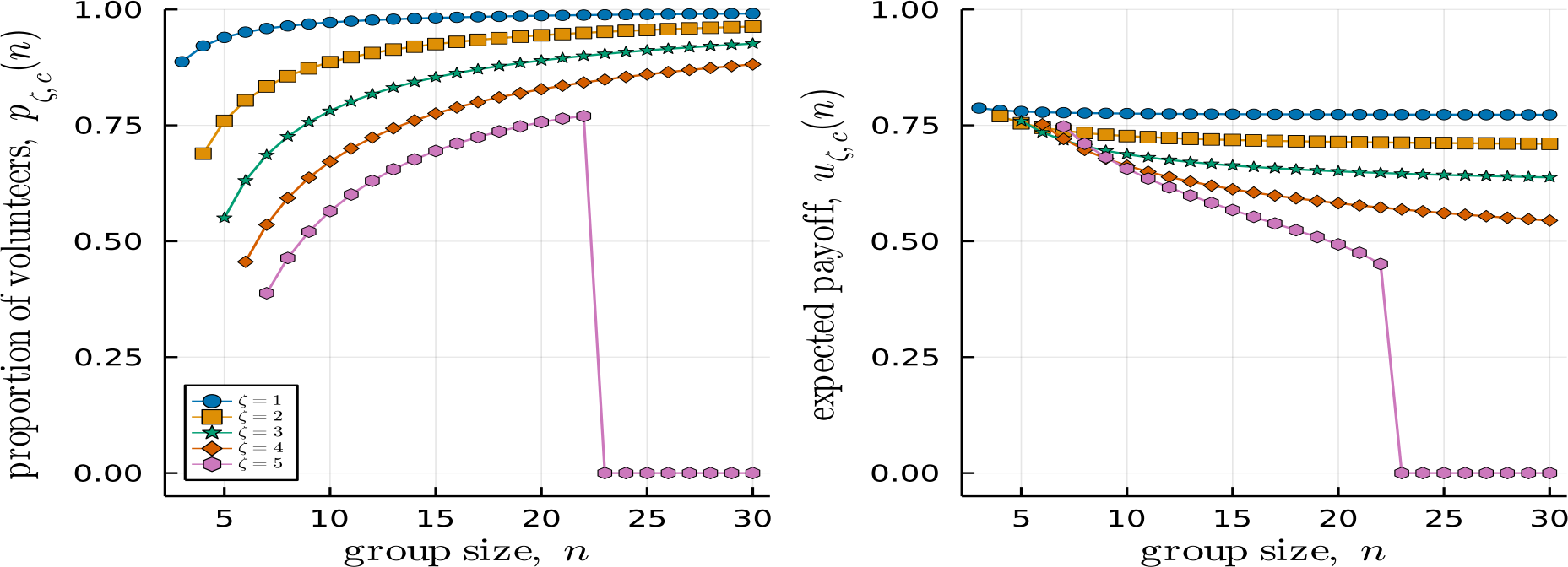
Proportion of volunteers *p*_*ζ,c*_(*n*) (*left*), and expected payoff *u*_*ζ,c*_(*n*) (*right*) as functions of group size for *ζ* ∈{1, 2, 3, 4, 5}, *c* = 0.2, and *n*∈ { *ζ* + 2, …, 30 }. For *ζ* ≤ 3 the inequality 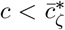 holds, so that the proportion of volunteers is strictly increasing (Proposition 1) and the expected payoff is strictly decreasing (Proposition 2) in group size. Even though 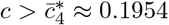 holds, for *ζ* = 4 the same monotonicity properties hold over the range of group sizes illustrated here because the critical group size is 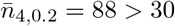. For *ζ* = 5 both the proportion of volunteers and the expected payoff drop to zero at the critical group size 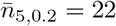.

**Figure 5.**
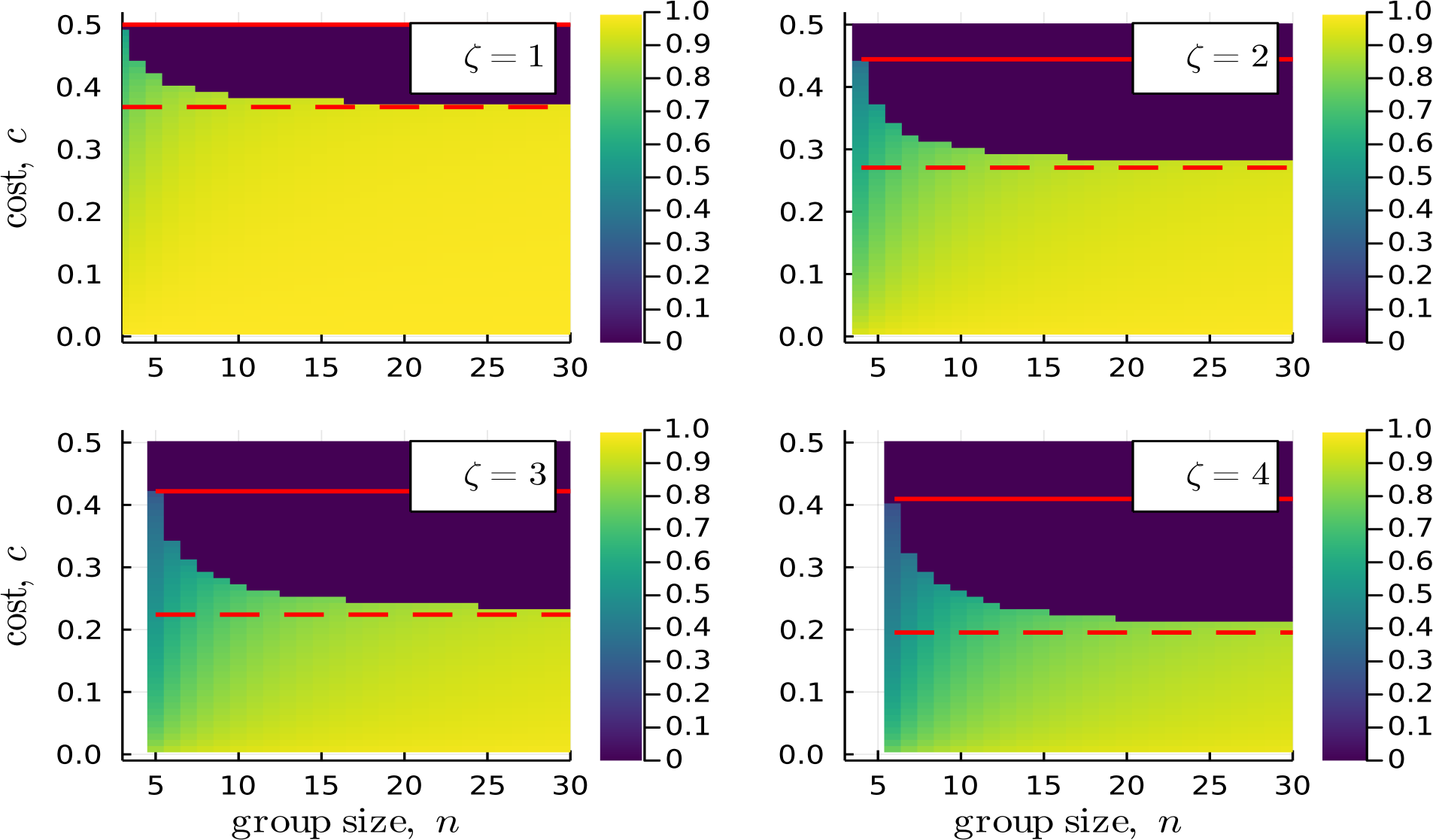
Proportion of volunteers *p*_*ζ,c*_(*n*) for *ζ* ∈ {1, 2, 3, 4}, *c* ∈ {0.01, 0.02, …, 0.5}, and *n* ∈ { *ζ* + 2, …, 30 }. The maximum and limit critical costs, 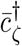 and 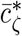, are shown respectively as *red solid* and *red dashed* lines. As proven in Proposition 1, for costs between these two critical costs, the proportion of volunteers is first strictly increasing and then drops to zero. For costs below 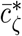(*red dashed line*), the proportion of volunteers is strictly increasing.

#### Proposition 1.

*For any ζ and* 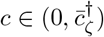, *the proportion of volunteers p*_*ζ,c*_(*n*) *is either strictly increasing or unimodal in group size n. More precisely:*

1. *If* 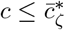, *then p*_*ζ,c*_(*n*) *is strictly increasing in n*.
2. *If* 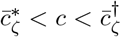, *then p*_*ζ,c*_(*n*) *is strictly increasing in n for* 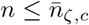*and equal to zero for* 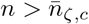.

This result demonstrates a positive group-size effect on the evolution of cooperation for the shirker’s dilemma. For sufficiently low costs (i.e., 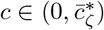) the proportion of volunteers at the minimal rest point is strictly increasing in group size, so that the larger the group size the larger the proportion of volunteers in the population at equilibrium. For intermediate costs (i.e., 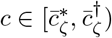) the proportion of volunteers at the minimal rest point is strictly increasing with group size up to the critical group size (for group sizes 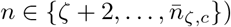before falling to zero thereafter (for group sizes 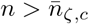). It follows that, from the perspective of maximizing the proportion of volunteers at the minimal equilibrium, the optimal group size is either intermediate and equal to the critical group size (for intermediate costs) or infinite (for low costs).

This said, note that Lemma 3 also demonstrates a negative group-size effect when looking into the comparative statics of the basin of attraction of the interior rest point 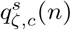 sustaining non-zero volunteering at equilibrium. Since the size of this basin of attraction is determined by the proportion of shirkers at the unstable rest point 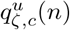 and this proportion decreases with group size, it follows that the basin of attraction of 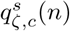 decreases (and the basin of attraction of full shirking increases) with group size (see Fig. 6 for an illustration). Overall, the proportion of volunteers at equilibrium can increase with increasing group size, but such an increase is accompanied by a decrease in its basin of attraction.

**Figure 6.**
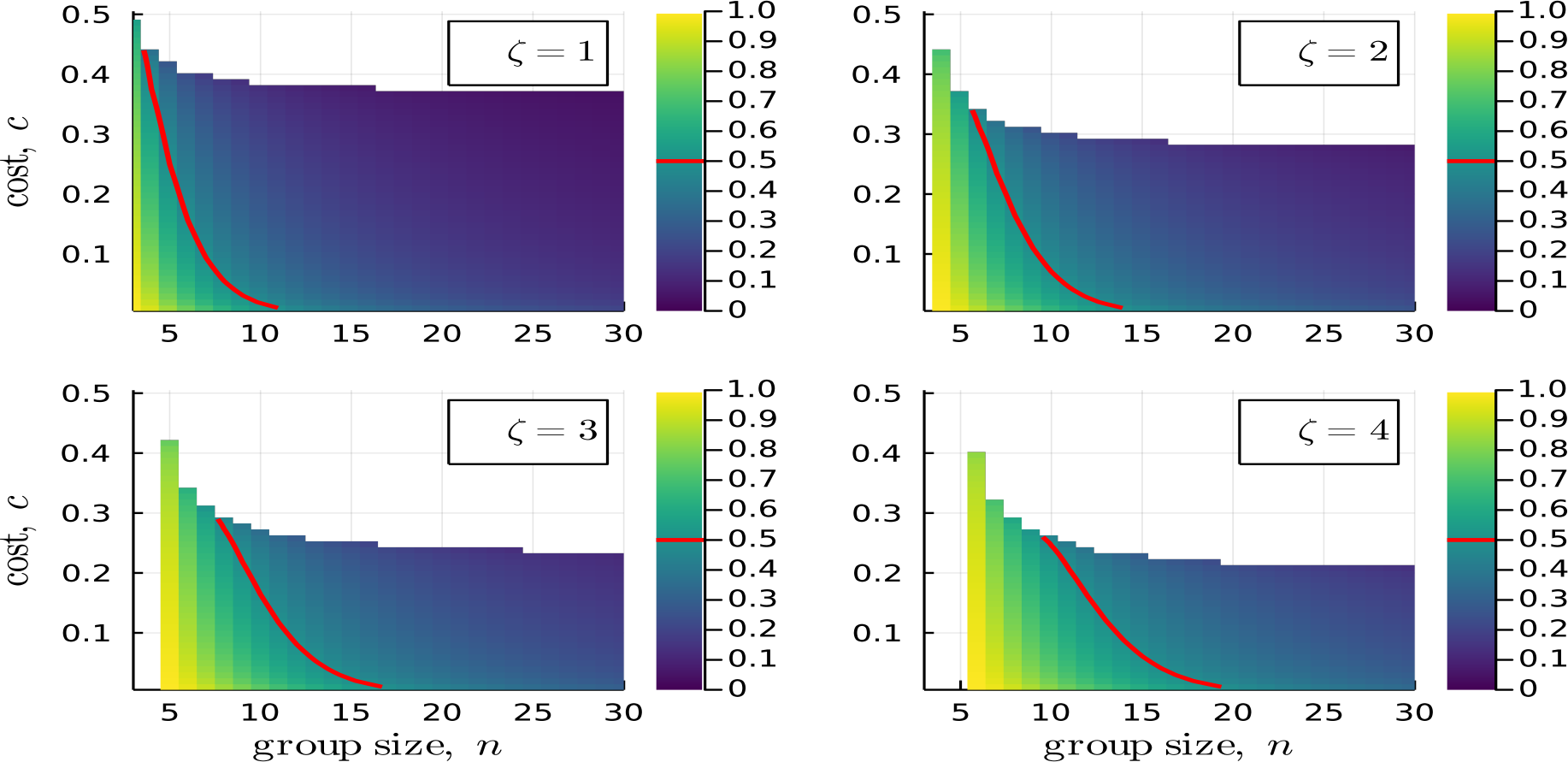
Location of the unstable rest point 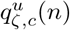, or equivalently, the size of the basin of attraction of the stable rest point 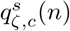 for *ζ* ∈ {1, 2, 3, 4}, *c* ∈ {0.01, 0.02, …, 0.5}, and *n* ∈ {*ζ* + 2, …, 30}. The *red solid line* represents the contour line at which 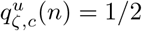 holds. To the left of this line, 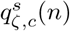 has the largest basin of attraction. To the right of this line, full shirking has the largest basin of attraction. As indicated in the main text, 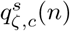 decreases with group size *n* for fixed threshold of shirkers *ζ* and cost of volunteering *c*.

### 3.3 The effect of group size on the expected payoff

Next, we address the effect of group size on the expected payoff. The following result shows that the group-size effect on the expected payoff at equilibrium is negative (see Appendix A.3 for a proof, and the right panel of Fig. 4 and Fig. 7 for an illustration).

**Figure 7.**
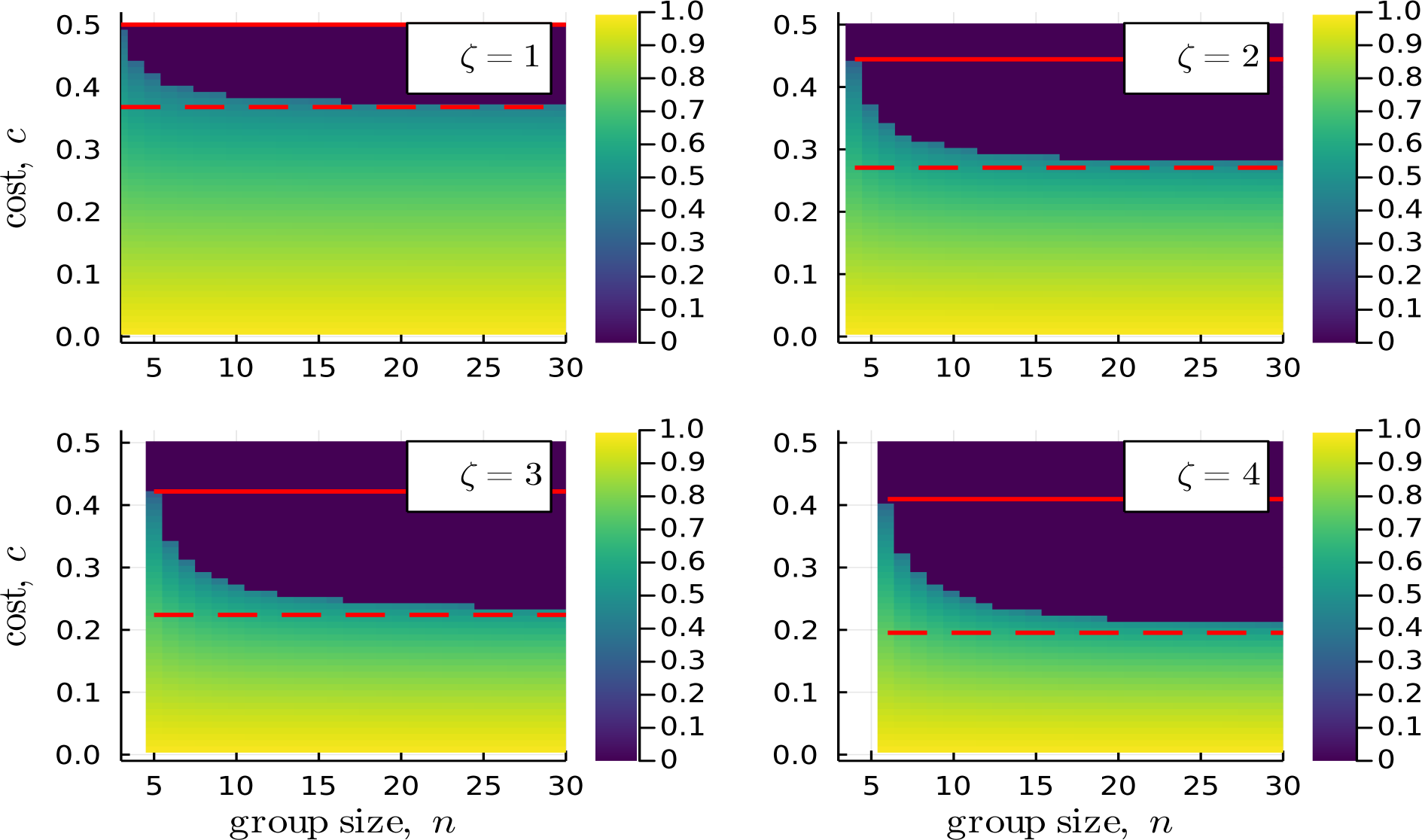
Expected payoff *u*_*ζ,c*_(*n*) for *ζ* ∈ {1, 2, 3, 4}, *c* ∈ {0.01, 0.02, …, 0.5}, and *n* ∈ {*ζ* + 2, …, 30}. The maximum and limit critical costs, 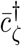 and 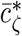, are shown respectively as *red solid* and *red dashed* lines. As proven in Proposition 2, for costs between these two critical costs, the expected payoff is strictly decreasing and drops to zero for finite but sufficiently large group sizes. For costs below 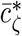(*red dashed line*), the expected payoff is strictly decreasing.

#### Proposition 2.

*For any ζ and* 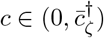, *the expected payoff u*_*ζ,c*_(*n*) *is decreasing in group size n. More precisely:*

1. *If* 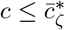, *then u*_*ζ,c*_(*n*) *is strictly decreasing in n*.
2. *If* 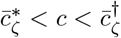, *then u*_*ζ,c*_(*n*) *is strictly decreasing in n for* 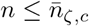*and equal to zero for* 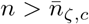.

This result contrasts with the positive group-size effect given in Proposition 1 when considering the proportion of volunteers. Together, Propositions 1 and 2 indicate that although there can be less shirking at equilibrium as the group size increases, the increased cooperation rate is not enough to increase (via a concomitant increase in the success probability, see Eq.(14)) the expected payoff, and hence the fitness, of individuals in the population.

### 3.4 The effect of group size on the success probability

Consider an increase in group size from *n* to *n* + 1. If a stable interior rest point exists for both of these group sizes, which from Lemma 2 will be the case if and only if 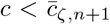, Propositions 1 and 2 give definite answers to the question of how such an increase in group size affects the proportion of volunteers and expected payoff at the stable interior rest point: the proportion of volunteers increases, whereas the expected payoff decreases. In contrast, the effect of such an increase in group size on the success probability at a stable interior rest point cannot be unambiguously signed.

To see why this is the case, it is instructive to use Eq.(14) to rewrite the success probability *ϕ*_*ζ,c*_(*n*) as

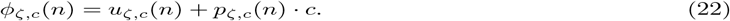

It is then apparent that the group size effect on the success probability is the sum of two effects pointing in opposite directions: an increase in group size from *n* to *n* + 1 reduces the expected payoff *u*_*ζ,c*_(*n*), but at the same time increases the proportion of volunteers *p*_*ζ,c*_(*n*). Whether the sum of these two effects results in an increase or a decrease in the success probability depends on their relative strength. In principle, it could be possible that one of the two effects is always stronger than the other, thereby allowing us to sign the group size effect on the success probability. However, the following proposition shows that this is not so by demonstrating that whether an increase in group size results in an increase or a decrease of the success probability depends on the cost parameter (the proof is in Appendix A.4).

#### Proposition 3.

*Suppose* 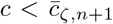*holds. Then the success probabilities for group size n and n* + 1 *satisfy:*

1. *ϕ*_*ζ,c*_*(n + 1) < ϕ*_*ζ,c*_*(n) for cost c sufficiently close to* 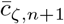.
2. *ϕ*_*ζ,c*_*(n + 1) > ϕ*_*ζ,c*_*(n) for cost c sufficiently close to zero*.

Proposition 3 indicates a negative (resp. positive) group size effect on the success probability for relative large (resp. small) costs. We can offer some intuition for the first of these results: When costs are close to but below 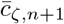, then (as can be seen from Fig. 3) the positive group-size effect on the proportion of volunteers at the stable interior rest point must be very small, implying that the change in the second term on the right side of Eq.(22) is very small, too. At the same time, when the increase in the proportion of volunteers is very small, then an increase in group size will have a significant negative effect on the expected payoff. Hence, the overall effect on the success probability must be positive. No such simple intuition is available for the second result. Indeed, the formal proof shows that the result hinges crucially on the relative rate at which the proportions of shirkers at the stable interior rest points for group sizes *n* and *n* + 1 converge to zero when the cost approaches zero.

The left panel of Fig. 8 illustrates Proposition 3 by considering three different costs. For the smallest of these (*c* = 0.305), the success probability for *ζ* = 1 increases with group size for all group sizes in {3, …, 10}, whereas for the largest of these (*c* = 0.31) the success probability decreases with group size.These cost levels are thus sufficiently small (resp. sufficiently large) for the results in Proposition 3 to be applicable. This is not the case for the intermediate cost level (*c* = 0.3075) for which the left panel of Fig. 8 illustrates that there are cases, not covered by Proposition 3, in which the success probability at an interior stable rest point can be first increasing and then decreasing.

**Figure 8.**
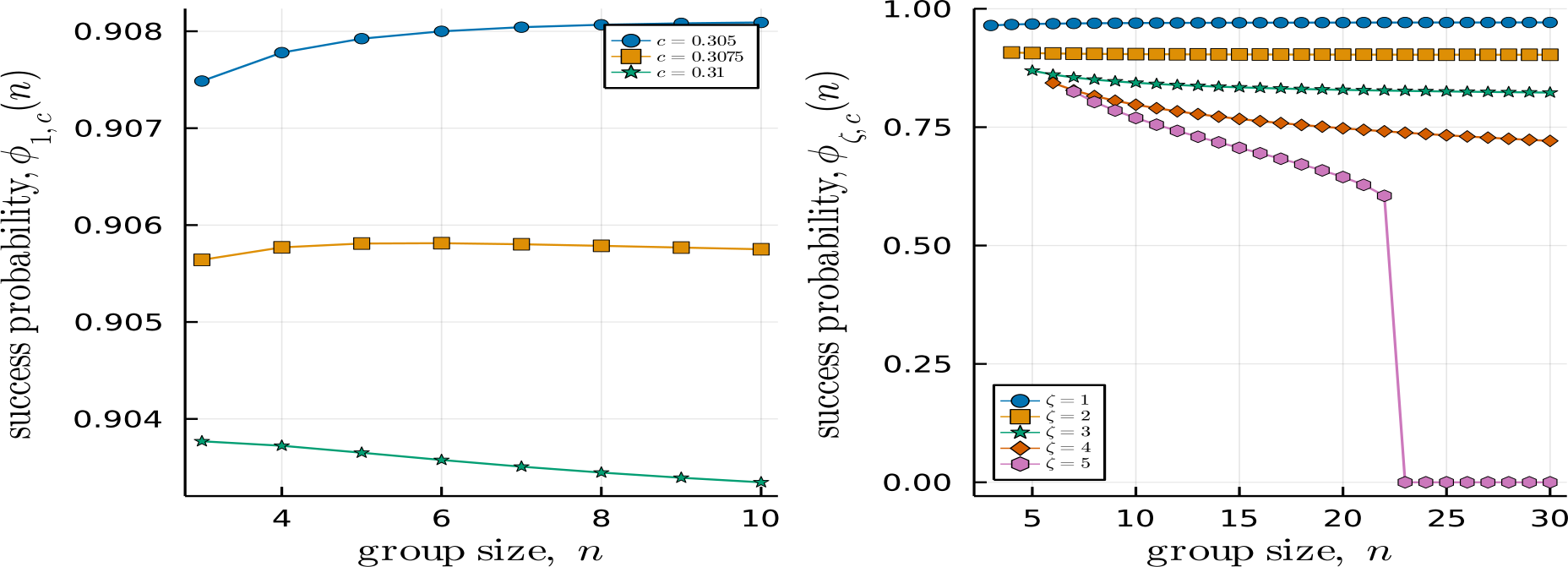
Success probability as a function of group size. *Left:* success probability *ϕ*_*ζ,c*_ for *ζ* = 1, three different values of cost *c*, and *n*∈ { 3, …, 10 }. For this range of group sizes, *ϕ*_1,*c*_ is strictly increasing for *c* = 0.305, unimodal (with a local maximum at *n* = 6) for *c* = 0.3075, and strictly decreasing for *c* = 0.31. *Right* : success probability *ϕ*_*ζ,c*_(*n*) for *ζ* ∈ {1, 2, 3, 4, 5}, *c* = 0.2, and *n* ∈ {*ζ* + 2, …, 30}. For *ζ* = 5, the success probability falls to zero at the critical group size 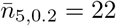.

We note that the success probability is high and does not change very much with group size for the three cost parameters considered in the left panel of Fig. 8,. The right panel of Fig. 8 illustrates a similar behavior of the success probability for low values of the shirker threshold (*ζ* ∈ {1, 2, 3}), whereas for higher values (*ζ* ∈ {4, 5}) there is a more pronounced effect of group size on the shirking probability. Fig. 9 illustrates the dependence of the success probability on the cost of volunteering and the group size for *ζ* ≤ 4 and suggests that, in most cases, the success probability remains high and almost unchanged across group sizes. A similar effect can be seen in Fig. 7, which illustrates the dependency of the expected payoff on group size. This prompts us to look into the limits of these two quantities and of the volunteering probability when the group size tends to infinity. We do so in the following section.

**Figure 9.**
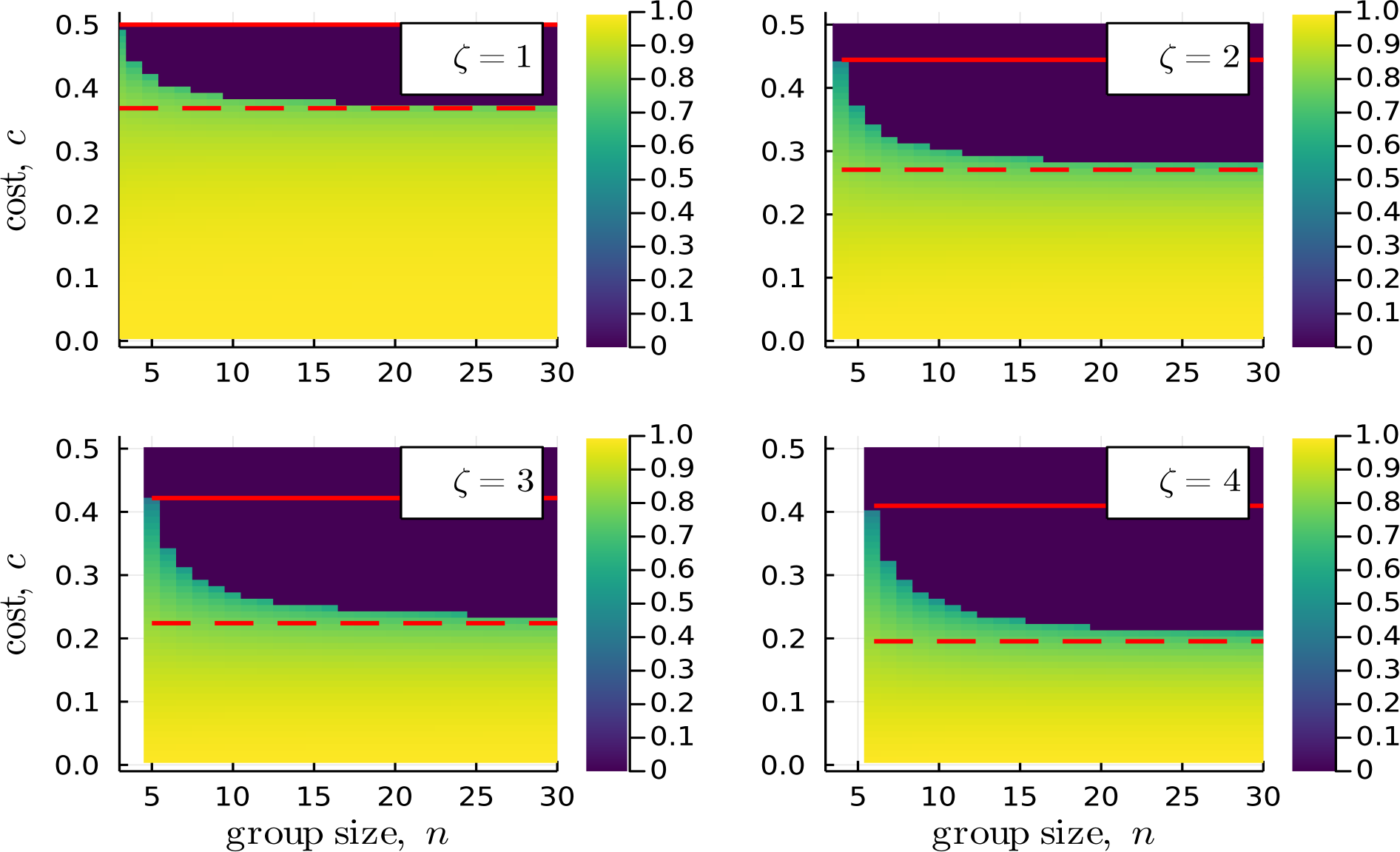
Success probability *ϕ*_*ζ,c*_(*n*) for *ζ* ∈ {1, 2, 3, 4}, *c* ∈ {0.01, 0.02, …, 0.5}, and *n* ∈ {*ζ* + 2, …, 30}. The maximum and limit critical costs, 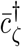 and 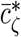, are shown respectively as *red solid* and *red dashed* lines.

### 3.5 The limit of infinitely large groups

From the results in Propositions 1 and 2 it is clear that the limits

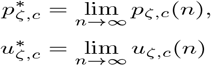

are well defined. Furthermore, it follows from Eq.(22) that

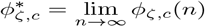

is given by

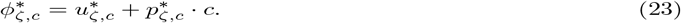

and thus also well-defined. In the following, we will determine the above three limits for the case where 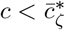 (as for 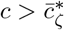 all limits are equal to zero and the knife-edge case 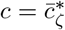 requires a case distinction without yielding additional insights).

We begin by observing that Lemma 1.3 implies *q*_*ζ,c*_(*n*) *< ζ/*(*n* − 1) for all *n*. It is then immediate that 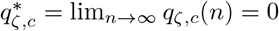holds. Using Eq.(11) we thus obtain 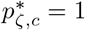, i.e., the proportion of volunteers in the minimal equilibrium converges to one. From Eq.(23) this in turn implies 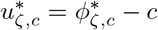, i.e., in the limit, the expected payoff differs from the success probability by the cost.

It remains to determine the limit of the success probability 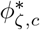. Even though the proportion of contributors converges to one, this limit is smaller than one. The reason is that the speed of convergence of *q*_*ζ,c*_(*n*) to zero is sufficiently slow to ensure that the expected number of shirkers

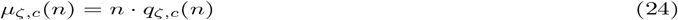

converges to a strictly positive limit. Indeed, we have (the proof is in Appendix A.5):

#### Lemma 4.

*Let* 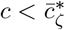. *Then*

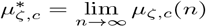

*is given by the unique solutions λ to*

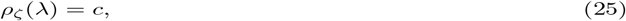

*in the interval* (0, *ζ*), *where ρ*_*k*_(*λ*) *denotes the probability mass function of a Poisson distribution with parameter λ (see Eq*. (17)).

Taking into account that lim_*n→∞*_ *q*_*ζ,c*_(*n*) = 0 implies that the expected number of shirkers (from the perspective of an outside observer) and other shirkers (from the perspective of a focal player) in a group coincide in the limit, Eq.(25) is the natural counterpart to the pivotality condition (8) when the number of shirkers among co-players follows a Poisson distribution with mean value *λ*, as it is the case in the limit for *n* → ∞. Further, just as Lemma 1 identifies the unique solution to the pivotality condition (8) in the interval (0, *ζ/*(*n* − 1)) as the (non-trivial) minimal rest point, Lemma 4 indicates that the limit value 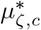 of the expected number of other shirkers in a group is the unique solution to Eq.(25) in the interval (0, *ζ*). See Fig. 10 for an illustration of the determination of 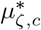 and of the following proposition, which completes our characterization of the limit (the proof is in Appendix A.5).

**Figure 10.**
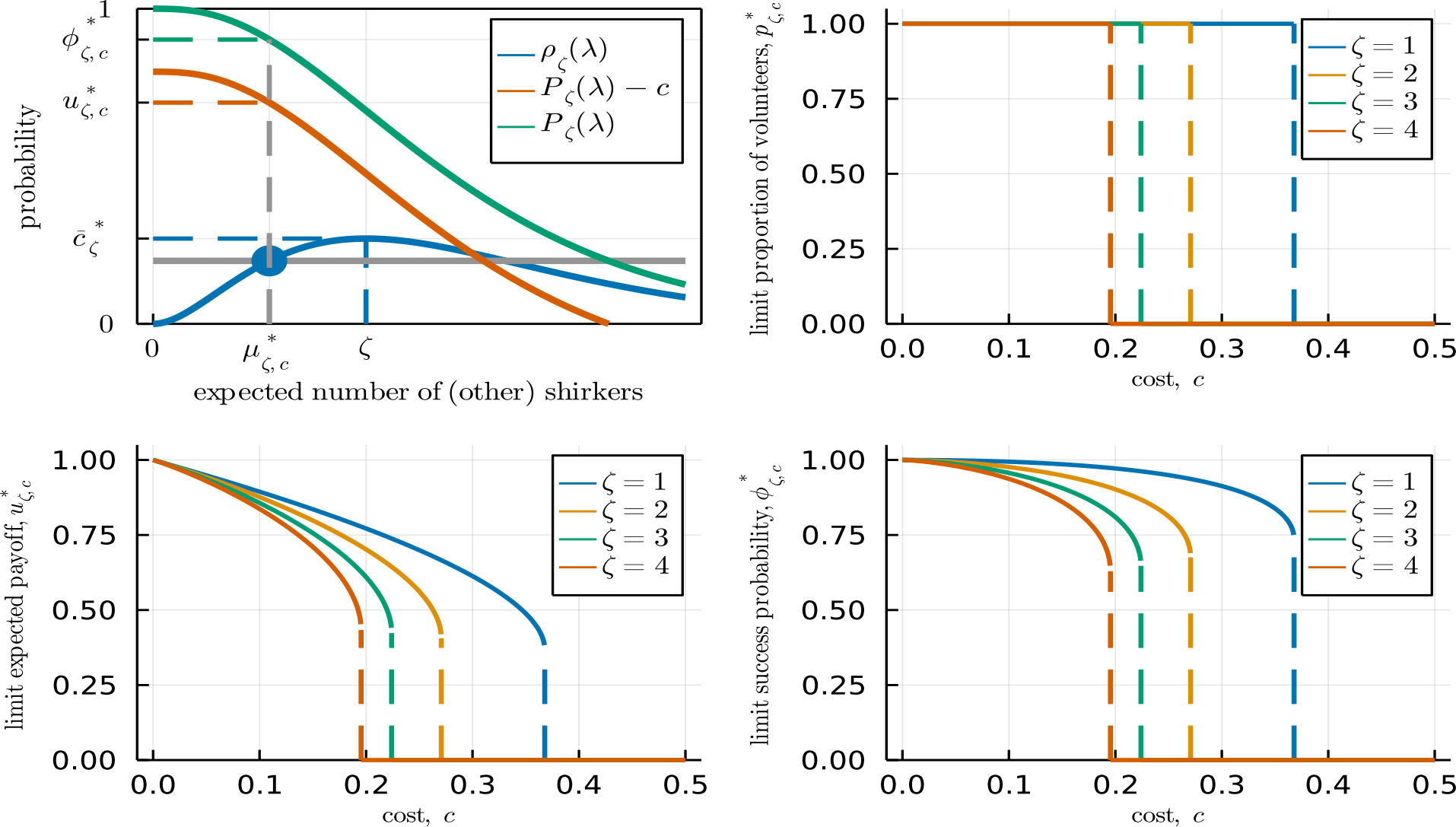
Illustration of the limit results in Lemma 4 and Proposition 4. *Top left:* The limit of the expected number of shirkers 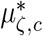 is given by the smaller of the two solutions to the pivotality condition *ρ*_*ζ*_(*λ*) = *c*. The success probability 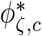 is the probability 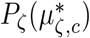 that there are at most *ζ* shirkers given that the number of shirkers follows a Poisson distribution with expected value 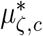. The expected payoff 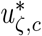 differs from 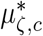 by the cost *c*. Here, *ζ* = 2 and *c* = 0.2. *Top right, bottom left, and bottom right:* Limit proportion of volunteers, 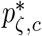, expected payoff 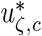, and success probability 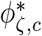as functions of cost *c*, for *ζ* ∈ {1, 2, 3, 4}.

#### Proposition 4.

*Let* 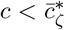. *Then, where*

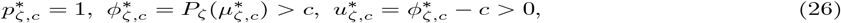

*where*

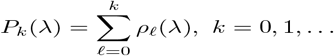

*is the cumulative distribution function of a Poisson distribution with parameter λ*.

Proposition 4 demonstrates that, for sufficiently low costs of volunteering, the proportion of volunteers at the minimal rest point converges to one as the group size tends to infinity. In such a limit, both the success probability and the expected payoff at equilibrium are positive values that, as illustrated in Fig. 10, can be relatively large for small values of the threshold number of shirkers *ζ*. In particular, even though the expected payoff at the minimal rest point decreases with group size (as demonstrated in Proposition 2), the expected payoff can still be substantially larger than the expected payoff at the full shirking equilibrium (which is zero). Likewise, the probability that collective action is successful can be relatively high. For instance, for a cost of volunteering *c* = 0.1 (so that the cost is equal to one-tenth of the benefit), the limiting success probabilities 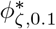 for *ζ* ∈ {1, 2, 3, 4} are always greater than 90% and given by 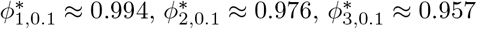, and 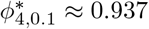. This said, note that a similar caveat concerning equilibrium selection as the one we pointed out for finite group sizes applies in the limit of infinitely large group sizes. In this case, it can be shown (by similar arguments demonstrating that 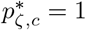 holds) that the size of the basin of attraction of the volunteering equilibrium 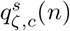 shrinks to zero.

## 4 Discussion

When the cost of volunteering is sufficiently small, the evolutionary dynamics of the shirker’s dilemma are characterized by two stable equilibria: a polymorphic equilibrium sustaining some cooperation (or a “volunteering” equilibrium) and a monomorphic equilibrium where everybody shirks (or a “full-shirking” equilibrium). We have investigated the group-size effects on the volunteering equilibrium and on a set of quantities related to it, namely (i) the proportion of volunteers, (ii) the probability that the public good is provided (or that the collective action is successful), and (iii) the expected payoff at such equilibrium, together with (iv) the size of the basin of attraction of the volunteering equilibrium. Our analysis reveals non-trivial comparative statics. On the one hand, we have found that, for all thresholds *ζ* ≥ 1 and sufficiently low costs, the proportion of volunteers at the volunteering equilibrium increases with group size, and converges to one in the limit of large group sizes. We also found that the probability that the public good is provided at equilibrium can increase with group size and converges to a positive value. On the other hand, the expected payoff, although positive, decreases with group size. Moreover, the basin of attraction of the volunteering equilibrium also decreases with group size and converges to zero in the limit of infinitely large group sizes. These findings are in contrast to those of the volunteer’s dilemma (Diekmann, 1985) and its generalization to threshold games in which a minimal number *θ* of ‘volunteers’ are needed (Palfrey and Rosenthal, 1984). For these games, both the proportion of volunteers and the overall probability that the public good is provided at equilibrium decrease with group size *n > θ* (Nöldeke and Peña, 2020). In the limit, the corresponding “volunteering” equilibrium of the volunteer’s dilemma tends to zero, and its basin of attraction tends to one. This result is the mirror image of what happens in the shirker’s dilemma analyzed in this paper.

We motivated the shirker’s dilemma in the Introduction with the examples of reproductive differentiation in *Dictyostelium discoideum* and the punishment of free-riders in multi-player social dilemmas. Another example is the sentinel behavior of Arabian babblers (*Argya squamiceps*): a territorial, cooperatively breeding species of songbirds (Zahavi, 1990) living in arid areas along the Great Rift Valley. During the day, group members take turns as sentinels on treetops, while the other group members forage on the ground or on low branches. When a sentinel spots an approaching raptor, it emits an alarm call. Foragers then have two options. First, they can flee to shelter inside a thicket, within which they are temporarily protected from the raptor but unable to follow its moves. Alternatively, they can fly up and join the sentinel on treetops in calling toward the raptor. The latter is the typical choice of foragers, constituting more than 80% of foragers’ reactions (Ostreiher and Heifetz, 2020). Moreover, even lone Arabian babblers devoid of territory (i.e., “floaters”) engage in calling from treetops towards raptors when they spot them (Ostreiher and Heifetz, 2017). This signal is costly in terms of both the energy expended on calling and the fatal risk in case they temporarily lose track of the maneuvering raptor. The raptor, on its part, should decide whether to continue hovering above the group until one of its members becomes less vigilant or, alternatively, to move on in search of other prey. Hiding group members are the least vigilant when they emerge out of the thicket, so moving on is better for the raptor than lurking for hiding birds only if the number of hiding birds (i.e., the number of “shirkers”) is small enough. Dissuading the raptor from attack is a public good enjoyed by all group members. This situation resembles more a shirker’s dilemma than a volunteer’s dilemma because the predator’s decision to hang around or to move away is likely to depend on the absolute number of non-vigilant individuals that it spots (i.e., the “shirkers”), rather than on the absolute number of vigilant individuals (i.e., the “volunteers”) that it is unlikely to catch. The fit of this example with the shirker’s dilemma model is not perfect, however, because if the public good is not provided and the raptor attacks the group, the shirker who ran to shelter and then comes out of it first is most likely to be targeted by the raptor, and thus bears a higher cost than the other group members, and in particular higher than those who joined the sentinel on treetops and did not shirk.

For another related example of a shirker’s dilemma, consider the mobbing of terrestrial predators (such as snakes) by groups of birds. When one group member reveals a snake dug in the sand and lurking for prey, that bird emits an alarm call, approaches the snake, and engages in bodily displays that make the bird look as big as possible. Noticing this, all or most group members then follow suit. In most cases, the primary and most important benefit of such mobbing is that it causes the predator to recognize that it has been detected, and to leave the location (Caro, 2005). Here, too, as in the case of a raptor approaching a group of Arabian babblers, the predator presumably moves away when it perceives that most of its potential prey are vigilant and that its outside options elsewhere might be more attractive.

The simple model we explored in this paper is a benchmark for more elaborate, relevant models. For instance, we modeled interactions in a well-mixed population and hence among unrelated individuals.

It would be of interest to see if our predictions are still supported when moving to spatially or family-structured populations where interactants are related, and where relatedness can be a function of group size (Lehmann and Rousset, 2010; Peña et al., 2015). Another dimension for possible extensions arises by considering games with continuous levels of effort and smooth benefits, rather than the binary-choice game with sharp thresholds that we analyzed in this paper. The pure equilibria of such continuous-action games do not predict the independent randomization characteristic of mixed-strategy equilibria of binary-action games, which received only partial empirical support in the context of the sentinel behavior of Arabian babblers (Heifetz et al., 2021). Further, such pure equilibria in continuous action games are not prone to the well-known peculiarity of mixed-strategy equilibria that individuals with higher costs or lower benefits cooperate with a higher probability at equilibrium (Diekmann, 1994). Considering continuous-action models would also allow us to explore if our result that shirking decreases with group size holds more generally or is a peculiarity of the binary-action models we have used here.

## Acknowledgements

J. Peña acknowledges funding from the French National Research Agency (ANR) under the Investments for the Future (Investissements d’Avenir) program (grant ANR-17-EURE-0010), from the Institute for Advanced Study (IAS) of the University of Amsterdam, and from the Max Planck Institute for Evolutionary Anthropology. A. Heifetz acknowledges funding from the Open University of Israel research grant 37142. The Julia code used for creating the figures of this paper is publicly available on GitHub (https://github.com/jorgeapenas/ShirkersDilemma).

## A Proofs

### A.1 Proof of Lemma 2

The equation *π*_*ζ,n*+1_(*q*) = *π*_*ζ,n*_(*q*) has a unique solution in the interval (0, 1), given by *q* = *ζ/n*. From the unimodality properties of the pivot probability *π*_*ζ,n*_(*q*) listed in Section 2.2, *ζ/*(*n* − 1) maximizes *π*_*ζ,n*_(*q*) over *q* ∈ (0, 1). Thus, recalling the definition of the critical cost 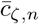 given in Eq.(9), we have 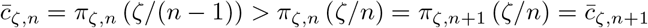. This proves that 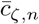 is strictly decreasing in *n* and thus maximized at the smallest possible value of *n*, which is *n* = *ζ* + 2. After setting *n* = *ζ* + 2 in (9) and simplifying this yields (15).

To prove the limit result, setting *m* = *n* − 1 in (7) and (9) and taking the limit we obtain

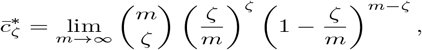

so that (16) follows from the Poisson approximation to the binomial distribution.

Finally, to prove that 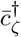 and 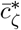 are decreasing in *ζ*, note that we can write 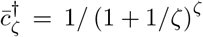 and 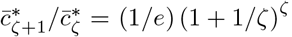. The function (1 + 1*/x*)^*x*^ increases with *x* for *x >* 0 (see, e.g., Hardy et al. 1952, Theorem 140) and its limit as *x* approaches infinity is *e*. Hence, 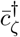 increases with *ζ* and 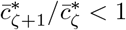 holds for all *ζ* ≥ 1.

### A.2 Proof of Lemma 3

For 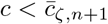 it follows from Lemma 1.3 that for group size *n* + 1 the replicator dynamics has two interior rest points satisfying 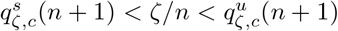. From Lemma 2 the inequality 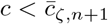 implies 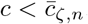. Hence, the replicator dynamics also has two interior rest points for group size *n*. Applying Lemma 1.3 again these rest points satisfy 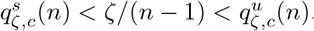. From Proposition 1 in Peña and Nöldeke, 2018 we have 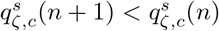 and 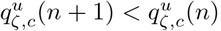. At the beginning of the proof of Lemma 2 we have established 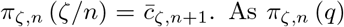. As *π*_*ζ,n*_ (*q*) is increasing in *q* for *q < ζ/*(*n* − 1) and 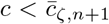 holds, the pivotality condition (8) implies the remaining inequality in (20), namely 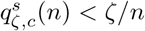.

### A.3 Proof of Proposition 2

It is immediate from Eq.(19) that *u*_*ζ,c*_(*n*) = 0 holds if *n* 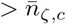. Further, it is also immediate from Eq.(19) that 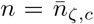 implies *u*_*ζ,c*_(*n*) *> u*_*ζ,c*_(*n* + 1) = 0. To prove the proposition it remains to consider the case 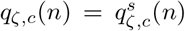 and 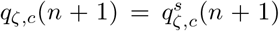. We may therefore assume throughout the following that *q*_*ζ,c*_(*n*) and *q*_*ζ,c*_(*n* + 1) satisfy the pivotality condition (8) and, from Lemma 3, satisfy 0 *< q*_*ζ,c*_(*n* + 1) *< q*_*ζ,c*_(*n*) *< ζ/n*.

Showing that *u*_*ζ,c*_(*n* + 1) *< u*_*ζ,c*_(*n*) holds is equivalent to showing that 1 − *u*_*ζ,c*_(*n* + 1) *>* 1 − *u*_*ζ,c*_(*n*) holds. From (2) and (13) this inequality is in turn equivalent to

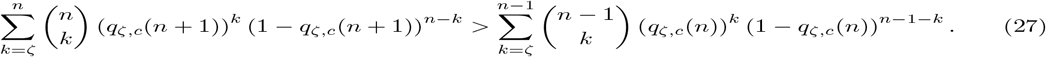

Begin by noting that (27) holds if the inequality

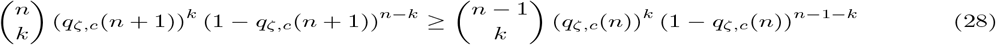

holds for all *k* ∈ {*ζ*, …, *n* − 1}, This is so because the last summand on the left side of (27), that is, (*q*_*ζ,c*_(*n* + 1))^*n*^, is strictly positive.

Let

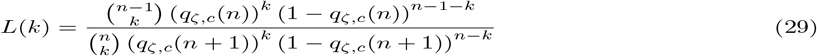

denote the likelihood ratio for having, among the co-players of a focal individual, exactly *k* shirkers at the minimal rest point with group sizes *n* and *n* + 1. Obviously, (28) is equivalent to the claim that *L*(*k*) ≤ 1 holds for *k* ∈ {*ζ*, …, *n* − 1}.

Because both *q*_*ζ,c*_(*n*) and *q*_*ζ,c*_(*n* + 1) satisfy the pivotality condition (8), we know that for *k* = *ζ* the terms on the left side and the right side of (28) are both equal to *c*. This shows *L*(*ζ*) = 1. From (29) we have *L*(*k* + 1) = *r*(*k*)*L*(*k*), where

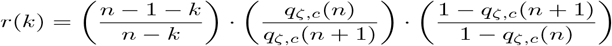

is strictly decreasing in *k*. Hence, provided that we can establish *L*(*ζ* + 1) ≤ 1 or, equivalently (from *L*(*ζ*) = 1), *r*(*ζ*) ≤ 1, our proof is finished (as *r*(*ζ*) ≤ 1 implies *r*(*k*) *<* 1 for all *k > ζ*, and thus *L*(*k*) *<* 1 for all *k > ζ* + 1).

We first demonstrate *r*(*ζ*) ≤ 1 for cost sufficiently close to (but below) the threshold 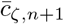 at which the stable interior rest point 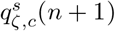 disappears. Straightforward calculations (see the beginning of the Proof of Lemma 2) show that

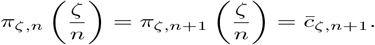

It then follows from Lemma 3 that both both *q*_*ζ,c*_(*n* + 1) and *q*_*ζ,c*_(*n*) converge to *ζ/n* as *c* converges to 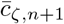from below. When *c* converges to 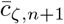 from below, *r*(*ζ*) thus converges to (*n* − 1 − *ζ*)*/*(*n* − *ζ*) *<* 1, implying that *r*(*ζ*) *<* 1 holds for all sufficiently high 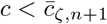.

Now suppose there exists some 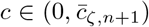 such that *L*(*ζ* + 1) *>* 1 holds. As *q*_*ζ,c*_(*n*) and *q*_*ζ,c*_(*n* + 1) are both continuous in *c* and the likelihood ratio (29) is also continuous in these probabilities it follows that there exists 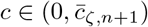 such that *L*(*ζ* + 1) = 1 holds. Fix such *c*. From *L*(*ζ*) = 1, we then have *r*(*ζ*) = 1. Because *r*(*k*) is strictly decreasing in *k*, we then have *r*(*k*) *>* 1 for all *k < ζ* and *r*(*k*) *<* 1 for all *k > ζ*. This implies *L*(*k*) *<* 1 for all *k > ζ* + 1 and also *L*(*k*) *<* 1 for all *k < ζ*. But this is impossible because the probabilities for obtaining *k* = 0, … *n* + 1 shirkers need to sum to one for both group sizes. We conclude that *L*(*ζ* + 1) ≤ 1 holds for all 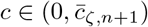, thus finishing the proof.

### A.4 Proof of Proposition 3

We have already noted in the proof of Proposition 2 that *q*_*ζ,c*_(*n* + 1) and *q*_*ζ,c*_(*n*) both converge to *ζ/n* as *c* converges to 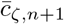 from below. From (12) the corresponding limit values of the success probabilities *ϕ*_*ζ,c*_(*n*) and *ϕ*_*ζ,c*_(*n* + 1) when 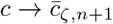 are given by Π_*ζ,n*_(*ζ/n*) for group size *n* and by Π_*ζ,n*+1_(*ζ/n*) for group size *n* + 1. Observing that Π_*ζ,n*+1_(*q*) *<* Π_*ζ,n*_(*q*) holds for all *q* ∈ (0, 1), it follows that *ϕ*_*ζ,c*_(*n* + 1) *< ϕ*_*ζ,c*_(*n*) holds for *c* sufficiently close to but smaller than 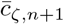.

The argument demonstrating the inequality *ϕ*_*ζ,c*_(*n* + 1) *> ϕ*_*ζ,c*_(*n*) for *c* sufficiently close to 0 requires a detailed investigation of the limit behavior of these success probabilities when *c* → 0. We thus proceed in a number of steps.

First, in the limit when *c* tends to zero both *q*_*ζ,c*_(*n*) and *q*_*ζ,c*_(*n* + 1) converge to zero, i.e.,

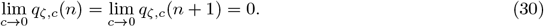

This is immediate from the properties of the pivot probabilities we noted in Section 2.3 and the fact (cf. Eq.(20) in the statement of Lemma 3) that the stable interior rest points *q*_*ζ,c*_(*n*) and *q*_*ζ,c*_(*n* + 1) lie in (0, *ζ/n*) for all 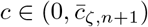 (and thus cannot converge to 1).

Second, it is immediate from (30) and the fact that the success probability is equal to 1 when the proportion of shirkers in the population is zero that

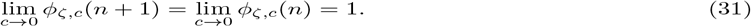

Hence, a sufficient condition for the inequality *ϕ*_*ζ,c*_(*n* + 1) *> ϕ*_*ζ,c*_(*n*) to hold for sufficiently small *c* is that the inequality

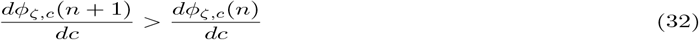

holds for all sufficiently small *c*.

Third, using (12) and the chain rule, we have

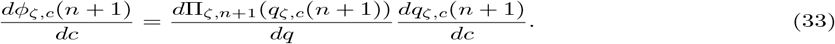

Either by direct calculation or by applying the derivative rule for polynomials in Bernstein form (see, e.g., Peña et al. 2014, Eq. 5), we find

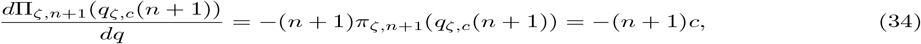

where the last equality uses the pivotality condition (8). Using (8) once more, we can apply the implicit function theorem to determine

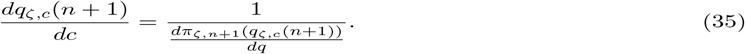

Combining these calculations by substituting (34) and (35) into (33), we have

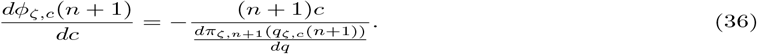

Similarly, we obtain

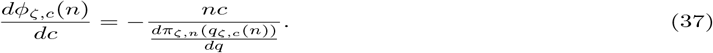

Substituting (36) and (37) into (32), and rearranging, it follows that condition (32) is equivalent to

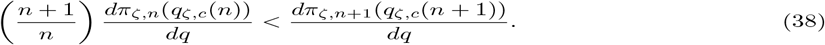

Fourth, we can again use either direct calculation or apply the derivative rule for polynomials in Bernstein form to the definition of the pivot probabilities in (7) to determine the two derivatives appearing in (38).

This yields

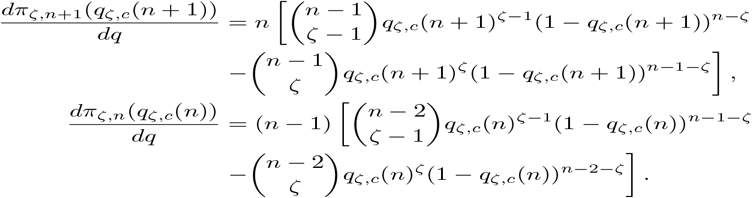

Using these expressions, we can rewrite (38) as

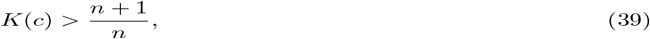

where

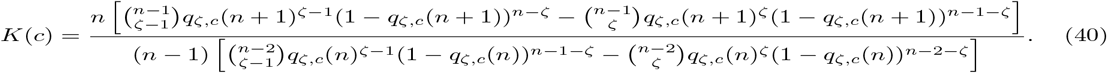

Fifth, we have

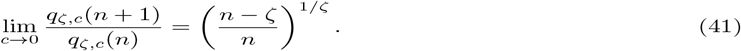

To see this, observe that from the pivotality condition (8) we have *π*_*ζ,n*+1_(*q*_*ζ,c*_(*n* + 1)) = *π*_*ζ,n*_(*q*_*ζ,c*_(*n*)) for all 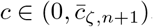. Substituting from the definition of the pivot probabilities in (7) it follows that

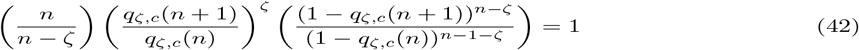

holds for all 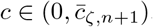. Clearly, the equality in (42) is preserved in the limit when *c* → 0. Using (30) to conclude that the third ratio on the left side of (42) converges to 1, we thus have

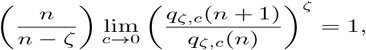

implying (41).

Sixth, making use of (30) and (41) it is straightforward to show that

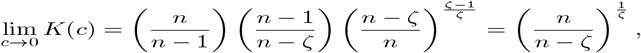

where *K*(*c*) had been defined in (40). It follows that (39) holds for all sufficiently small *c* if

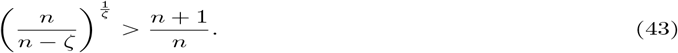

Observe that for *ζ* = 1 the inequality in (43) reduces to *n*^2^ *>* (*n* + 1)(*n* − 1), which holds for all *n*.

Seventh, to finish the proof it remains to establish (43) for 1 *< ζ < n* − 1. As we have already seen that (43) holds for *ζ* = 1 it suffices to argue that for any given *n* ≥ 3 the left side of (43) is increasing in *ζ* over the relevant range. Using the change of variable *x* = (*n* − *ζ*)*/ζ* this follows from the fact that (1 + 1*/x*)^*x*+1^ is decreasing in *x* for *x >* 0 (see, e.g., Hardy et al. 1952, last line on page 102).

### A.5 Proofs of Lemma 4 and Proposition 4

Let

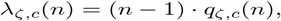

and

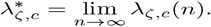

Taking into account that Lemma 1.3 implies *λ*_*ζ,c*_(*n*) *< ζ* for all *n*, arguments that are otherwise identical to the ones in the proof of Lemma 3 in Nöldeke and Peña, 2020 establish Lemma 4 provided that 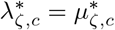 holds. As both *μ*_*ζ,c*_(*n*) − *λ*_*ζ,c*_(*n*) = *q*_*ζ,c*_(*n*) and lim_*n→∞*_ *q*_*ζ,c*_(*n*) = 0 hold, this is the case.

We have already noted in the text that the equalities 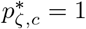 and 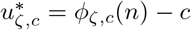 in the statement of Proposition 4 follow from Lemma 1.3. From a generalization of the classical Poisson approximation (see, e.g., Billingsley 1995, Theorem 23.2) 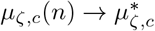 implies 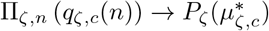. From (12) and the definition of 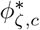 this implies 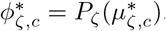, establishing the remaining equality in (26). An analogous argument, using the equality 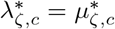 established in the above proof of Lemma 4 and Eq.(13), shows that 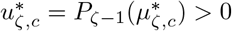, where the inequality follows from 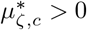. The proof is finished by observing that the inequality 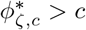 then follows from 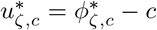.

